# Identifying key features of digital elements used during online science practicals

**DOI:** 10.1101/2023.03.02.530781

**Authors:** Vanda Janštová, Petr Novotný, Irena Chlebounová, Fina Guitart, Ester Forne, Montserrat Tortosa

## Abstract

As in everyday life, we use more digital elements as part of formal and informal education. To serve their educational purpose well, systematic research is desirable on identifying and measuring their characteristics. This study focuses on science practicals, which are complex and vary in organizational settings and specific arrangements, including usage of digital elements. We describe digital resources on which teaching online science practicals during COVID-19 forced lockdowns were built. We identified their key characteristics, as part of the Erasmus+ project ‘*My Home – My Science Lab*’, where science teachers in Slovakia, Czechia, Slovenia, France, and Spain shared web resources they used and would recommend and why. We recorded 89 inputs representing 50 unique web resources. Teachers preferred free web resources, mostly for knowledge revision, and newly discovered half of them due to forced distant teaching. The best evaluated resources were those supporting interaction (especially among peers), focused on teaching subjects and/or ICT, ready to use, and with a clear structure. The web resource most frequently mentioned and used in all countries was PhET (Interactive Simulations for Science and Math) providing free science principles simulations. Other characteristics mentioned in the literature (eg., supporting creativity and independent solving, connecting different levels of organization, authenticity, flexibility) were not that important for the overall rating.

## Introduction

As in other sectors of human activity, education in schools is subject to changes associated with the growing inclusion of digital elements used by both students and teachers. For decades, science education has been using digital elements such as videos and other visual material or domain-specific simulations, nowadays published mostly in the form of online activities. Online materials are easy to use in school and as homework, and are nowadays hardly seen as a technological novelty or innovative element of teaching, rather we consider them as a natural part of the educational process that corresponds to today’s educational reality.

The frequency of their use and the types of digital elements that teachers used, and the mode of use varied depending on the personality of the teacher, the technological equipment of the school, and also the school philosophy. Our focus is on the practice-oriented components of science teaching, by which we mean hands-on exercises, lab work (referred as *science practicals* in this study) which, at least in our experience, have been implemented online rather infrequently.

A fundamental change was caused by the COVID-19 pandemic and the transition of schools from conventional to online learning in many developed countries of the world, including Europe. This change has affected all teachers equally, regardless of their previous attitude towards digital technologies, creating a unique experimental and research space that has attracted the attention of many studies.

With the transition to online teaching, teachers had to modify science practicals to a form suitable for online teaching. The online form was then implemented even by teachers who were not technology experts. Such a necessity caused by external influence dramatically broadened the teachers’ experience of the online resources on offer, their possibilities, and ideas about the possibilities for use in their own teaching. As shown by (1), to adopt learning which includes (mobile) computing devices, it is necessary to go through several stages, where two are the critical ones: i) introduction and gaining practice, and ii) implementing the tools into lessons. All teachers in the European countries that we focus on have gone through both of these stages during distance teaching, including those who teach effectively, but have so far tended to avoid using technology in their teaching. As researchers, we thus see an opportunity to gain insight into the teachers’ evaluation of the digital elements, that is, to answer the question which digital elements characteristics are markers for teacher acceptance and use in teaching, suppressing the bias of technology acceptance (2, 3). It has been shown that the effectiveness of online teaching does not appear to be technology dependent, and teachers who taught more effectively face-to-face were more likely to teach more effectively also in the online setting (4). At the same time, (5) argue that the digital elements should be kept even in face-to-face learning as supplementary support materials to enhance student understanding, or possibly as optional distance education resources for students unable to be present in the laboratory, and as training materials for teaching assistants. Therefore, the study of digital elements in education is a current topic as we need to better understand their role in the teaching and learning process.

### Theoretical background

#### Forced online teaching

When the COVID-19 pandemic started, teachers in European countries (and not only they) were quickly adapting their courses from face-to-face to online due to massive lockdowns. It was a great challenge for teachers and instructors to quickly change the format of their in-person courses to remote courses. This transition to technologies in education was driven by needs and affected all teachers non selectively (6). Because they had to react from day to day, they tend to focus on gaining general technological skills specific for the online instructions followed by content (re)transformation, and to pay less attention to the integration of the technology with pedagogy and content (7). Such integration is important for effective teaching and, in general, similarly to other teaching settings, while covering and combining cognitive, affective, behavioral, and social aspects through communication and collaboration to engage the learner (8). Based on trying and searching for solutions and reflection, recommendations for conducting online teaching were formulated in parallel by (9) and (10). Based on these two classifications, we identified the following relevant online practical teaching components which are used in this study. These include: i) *building relationships and community:* understood as interactions that build relationships both within school (among students and between students and teachers) and outside school (within family, between students, and nature, students and society); ii) *active learning:* e.g., choice in the path to the learning goal, independent solving; iii) *learner autonomy* and supporting creativity; and iv) *personalized learning process:* which values, respects, and accommodates learner differences by providing learning materials in a variety of formats and modalities and allows independent exploration.

As pointed out by (11) and (12), the online learning environments were often originally used in hybrid or even in-person settings, which could compensate for their weaknesses and still benefit from the positives, e.g., bigger flexibility (13). Therefore, when designing online learning environments, socio-emotional processes have not been the main focus. Such environments were typically designed with productivity in cognitive processes and task related activities; not social interactions in mind (11, 12). And whether or not the socio-emotional processes are overlooked, the online environment is specific, as can be seen on an example of Bloom’s taxonomy where additional verbs typical for the digital environment were suggested (14). As a consequence, reduced and impeded social interaction was among the most pressing concerns among students during the COVID-19 pandemic (15). Many authors agree that communication and social relationships are especially challenging in online settings (16, 17).

#### Science practicals

The science practicals have had an important role in science education and have been shown to be a challenging method even in face-to-face settings. Their implementation is often considered to be more demanding even in face-to-face teaching: preparation, safety standards, organisation, interpretation of unexpected results, and unpredictability. They are effective in making students manipulate objects according to a recipe (18, 19), typically offering a unique possibility of working in a setting of smaller class sizes, higher instructor-to-student ratios, and the diverse learning possibilities of laboratory classrooms. Practicals can be effective and develop conceptual understanding when it is ‘hands on’ and ‘minds on’, and when teachers explicitly guide students to link these two essential components of practical work (20, 21). Also, in addition to improving knowledge, practical work has been shown to improve attitudes toward the subject (22, 23), interestingly in all of the following settings: traditional (not using computers), computer-supported laboratory and computer simulation (22).

The interest of researchers in science practicals settings led to the formulation of the design principles of students’ resources. These principles were developed regardless forced online teaching by (24) and include the ones relevant for our study which we highlight: i) *safety* – training to work in safe settings, in the online environment understood as the possibility to safely study dangerous or inaccessible features/processes; ii) *authenticity* - being based on real word/research situations; iii) *flexibility* - provides variety of topics and difficulty levels.

#### Digital elements in science practicals

The typical resources used for face-to-face laboratories had to be altered (or even replaced by a new format) from the base for distant teaching, and instructions started to emerge shortly after the first covid-19 forced lockdown. (25) analyzed different types of adjustments made to laboratory curricula due to COVID-19 pandemics (e.g., different types of experiments, simulation, pre collected data analysis, planning, reviews, etc.) and the reported immediate effect on students. The lack of hands-on laboratory experience was reported to be counterproductive to some types of learning and participation, such as understanding procedures, analyzing non-ideal data, evaluating the trustworthiness of the data, troubleshooting, making evidence-based conclusions, proposing experiments, or predicting results (25, 26). However, some goals could be achieved remotely (25) with the recommended setting, e.g. including peers in the learning formulated by (9,10,24). Specific recommendations for science practicals were introduced by (25–27) and are in concordance with the general rules. (28) point out that the fact that most of the students nowadays could use their smartphones for data recording, taking pictures, and connection to online apps such as virtual simulations helped to reach the science practicals goals.

For online science practicals, several *digital elements* have been described. I) *Investigation*, experiments, e.g., in kitchen settings which can be combined with online tools, for biology possibly including individual nature visits, e.g., sample collection outdoors like making a (digital) herbarium, includes remote experiments (25, 29). II) *Virtual reality*, which provides realistic interactions with 3D computer generated learning environments, or digital lab environments which are entirely in a virtual interface (e.g., Labster, Beyond Labz). Virtual laboratories can provide a safe learning environment accessible online, helping to better understand the experiment as a whole. The recommended setting is to use virtual laboratories before the real ones as a preparation for face-to-face laboratory experiments, as they lower stress level and anxiety and improve learning (5,30–32). And this is where the paradigm shift became apparent, where their role was seen as an adjunct to the face-to-face setting, but during forced online learning it became a separate form. During distant teaching, virtual laboratories were often substitutes for wet labs. III) *Computer simulations* have been shown to improve conceptual understanding of chemistry (33) and physics (34). The second study shows an example where students using computer simulations explicitly modeling electron flow understood the electricity concepts better and also could assemble the real circuit and describe how it worked better than students working in a real laboratory. IV) *Data analysis and computational science* (using computers to solve science problems) demonstrate the current approach in all science disciplines - solving, for example, molecule structure, molecular genetics (35) or enzyme kinetics (36). V) *Video-based learning* can be in many styles, from simple pre-recorded video experiments to real-time lab recordings (37).

For our work, we use a wider scope of these elements and also include some not-science specific components such as VI) *knowledge revision*, e.g. quiz which have been shown to significantly improve knowledge retention (38, 39); VII) *games* including a story line and rewards when students have to reach a goal either individually or in a team (40); VIII) *modeling and construction* when students learn by doing e.g. construction a measurement tool with Arduino followed by use of the mobile app ‘Arduino Science Journal’ (41); and IX) *virtual field* trip when the ‘visitors’ do not go to the actual site but ‘visit’ it through a virtual tour (42, 43). The types of elements mentioned demonstrate the difference between online and face-to-face teaching, where it is not simply a change of communication channel, but a qualitatively different type of work. The role of the digital elements in the online setting can be different from face-to-face settings and take on new functional roles, where, for example, a quiz is not only an activation and feedback tool but also a quick way of presenting the results of measurements or an assumption of the outcome of the intended observation. All these components contribute to the workflow of the online lesson and (26) suggest that they should be kept in face-to-face teaching, as they offer cost-efficient, environmentally friendly, and safe alternatives. They can be used as supplementary support materials to improve student understanding, or possibly as optional distance education resources for students unable to be present in the laboratory, and as training materials for teaching assistants (5).

New digital elements are continuously emerging, worthy of the interest of the general teaching public. Forced online learning has accelerated the ‘coevolution’ of digital elements developers and their users, clarifying the requirements for their functions and teachers’ demands for the inclusion of aspects (e.g., communication) that might have been neglected before. Therefore, we consider it important to be able to define their qualitative characteristics and to know which ones are decisive when teachers decide whether to adopt them in their future (face-to-face) practice. We start from the principles formulated by (9, 24) and combine them to collect data about digital elements used during online science practicals.

#### Aims

Our aim was to identify web resources that were adopted by science teachers during lockdown and incorporated into online science practicals; and to propose an interpretation of their shared characteristics which made them useful to teachers when planning effective online science education.

## Methods

We used an online survey to collect data about web resources from secondary school science teachers.

### Sample and sampling

Web resources were collected in 5 EU countries: Czechia, Slovakia, Slovenia, France and Spain from in-service science teachers using an online survey (GoogleForm) during the Erasmus+ project ‘*My Home - My Science Lab*’. Teachers were contacted by country coordinators based on their contacts and possibly also previous cooperation, and asked to share only activities they knew, had used and considered useful for online science teaching. Because the survey was anonymous and asked only for information about the reported web resources (and not about the teachers themselves), it was not necessary to seek approval from the ethics committee (Charles University, Faculty of Science). Teachers agreed to the use of the data provided by completing the survey with informed consent at the beginning.

### Instrument

Teachers provided the name of the web resource, hyperlink, overall rating of a web resource (scale 1-5, the more points, the better rating), information about synchronicity and subject (biology, chemistry, physics, geology, geography, STEM). The appropriate educational levels ISCED 1-3 were recorded together with information about availability free or after payment.

The pedagogical characteristics of the web resources and subsequently the digital resources were sampled in the following sections: 1) type of *digital element* (experiment, video, knowledge revision, augmented reality, simulations, etc.) with classification based on literature, see Table 1; 2) *pedagogical suitability*; 3) *interactions*; and 4) specific *added value* which are described below.

**Table 1.** Digital elements used in the survey. We provide a short definition and literature from which we draw.

| **Digital element** | **Definition** | **Literature** | **Acronym used in text**<br>*DE_learning digital element* |
| --- | --- | --- | --- |
| Experiment including remote experiment and demonstration | Students control variables, and have to observe and interpret the results | (25) | *DE_experiment* |
| Knowledge revision | Pre-prepared knowledge revision in a different form | (38) for quizz | *DE_revision* |
| Literature / data based analysis | Drawing conclusions from the literature, analyzing pre-collected sample data | (25,26) | *DE_literature* |
| Virtual field trip | Virtual tour / site visit | (42,43) | *DE_virtual trip* |
| Video based learning | Simple video recordings, narrative/voice-over lab recordings, and real-time delivery | (25,26) | *DE_video* |
| Simulation | Simulations of processes<br>focused on teaching of the<br>concepts | (25,44,45) | *DE_simulation* |
| Virtual / augmented<br>reality | Augmented reality (AR) and<br>Virtual Reality (VR) providing<br>realistic interactions with 3D<br>computer generated learning<br>environments | (26,31,46) | *DE_virtual reality* |
| Modeling,<br>constructing | learning by doing, e.g. devices<br>build with Arduino | (41) | *DE_modelling* |
| Games | Reaching a goal with an<br>identity created for the game,<br>can be played in teams | (40) | *DE_game* |
| Investigation | Inquiry; students investigate a<br>scientific phenomenon,<br>different levels (open,<br>structured, individual, in<br>groups) | (24) | *DE_investigation* |

In the next section of the survey, teachers reported the *pedagogical suitability* of the activity as the level of their agreement with statements describing the ease of use, suitability of the online tool for teaching science practicals and teaching ICT and other skills, and willingness to use the activity as is or after modification during online teaching or standard teaching on a 5-point Likert scale. Teachers rated the support for *interactions* among students, students and teachers, students and society, students and families, students and nature to assess the support in building relationships and community (using a 5-point Likert scale). Practicals are appreciated for specific *added values* as shown by the research; the extracted categories used for this study are summarized in Table 2. Except for the categories based on principles formulated mostly by (9, 24), we reported two more: providing worksheets and connecting different organizational levels. Providing worksheets to students is often part of science practicals (47), and ready-to-use worksheets are typically appreciated by teachers. Connecting different organizational levels is necessary to understand science and is described as an issue especially in biology (48, 49).

**Table 2.**
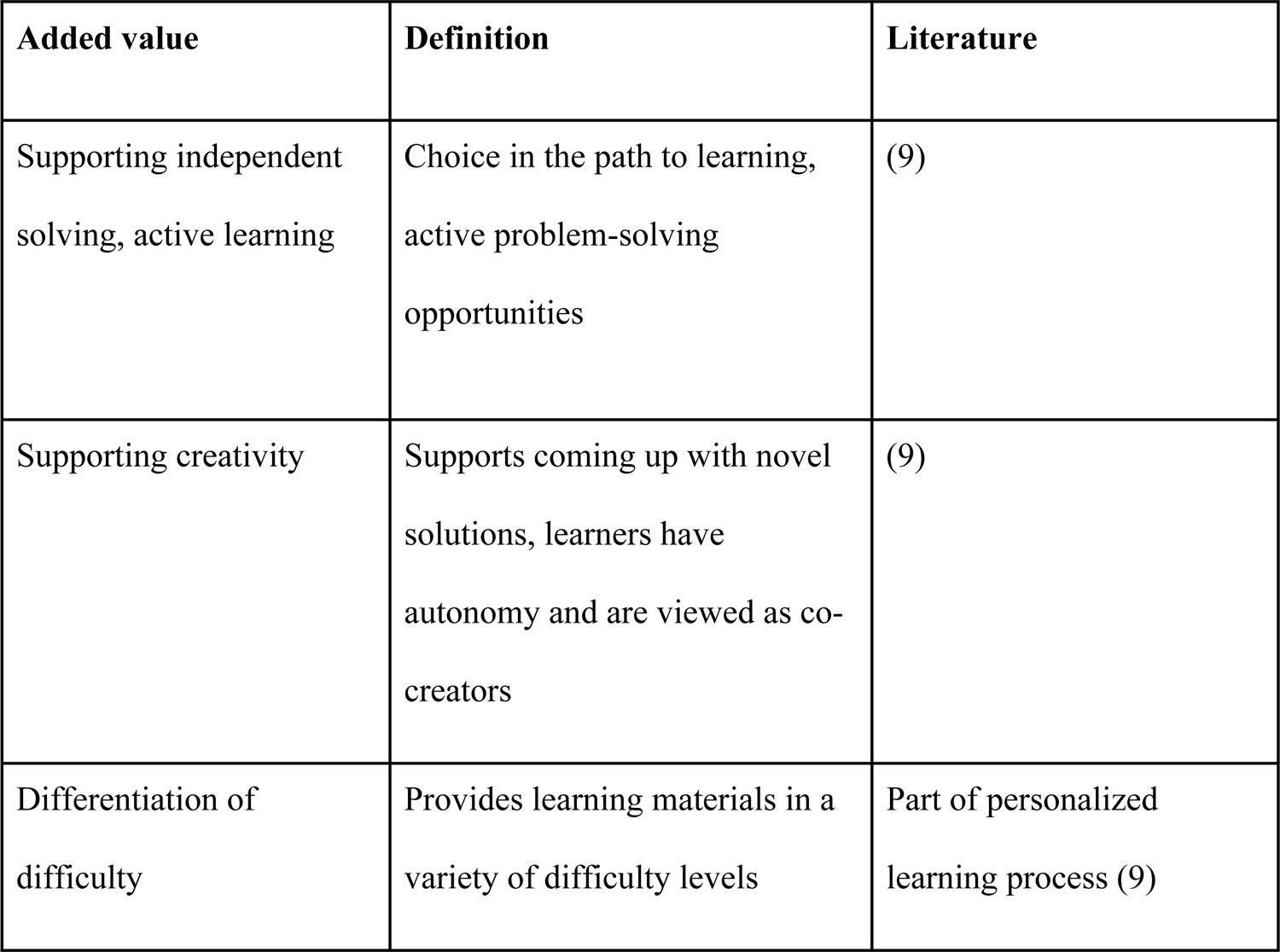
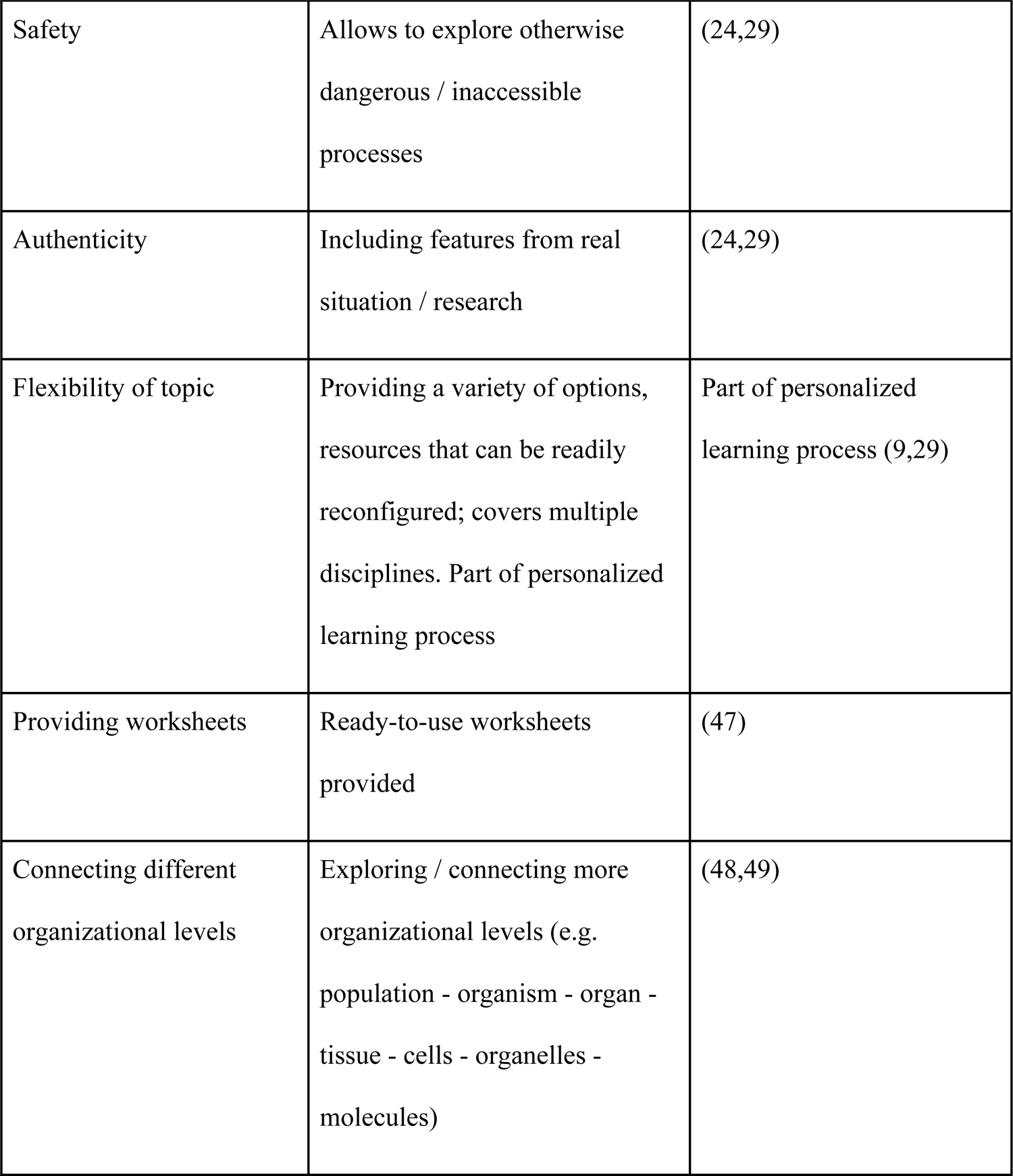
Web resource *added value* reported/studied overview.

| Added value | Definition | Literature |
| --- | --- | --- |
| Supporting independent solving, active learning | Choice in the path to learning, active problem-solving opportunities | (9) |
| Supporting creativity | Supports coming up with novel solutions, learners have autonomy and are viewed as co-creators | (9) |
| Differentiation of difficulty | Provides learning materials in a variety of difficulty levels | Part of personalized learning process (9) |
| Safety | Allows to explore otherwise dangerous / inaccessible processes | (24,29) |
| Authenticity | Including features from real situation / research | (24,29) |
| Flexibility of topic | Providing a variety of options, resources that can be readily reconfigured; covers multiple disciplines. Part of personalized learning process | Part of personalized learning process (9,29) |
| Providing worksheets | Ready-to-use worksheets provided | (47) |
| Connecting different organizational levels | Exploring / connecting more organizational levels (e.g. population - organism - organ - tissue - cells - organelles - molecules) | (48,49) |

The authors also additionally recorded whether the resource aimed at higher levels of cognitive processes of the revised Bloom taxonomy in factual and/or conceptual knowledge dimensions (50) or at ‘remembering and reproducing’. Among others, the key terms defined by (14) for the digital environment were used. The simultaneous assignment of the web resource to the knowledge revision and Bloom’s ‘remembering and reproducing’ categories was taken as a coding check.

### Data cleaning

The data was cleaned in Excel. The categories of digital elements which were not mentioned or were mentioned only once (*DE_virtual trip*, *DE_modelling*) were not included in the analysis. If the question allowed multiple answers, the format was transformed to dummy variables. All the answers were translated into English. In the case when a web resource contained one type of digital element (eg. simulation), but teachers reported more different digital elements’ types (eg. simulation, experiment, virtual experiment), the terminology was unified.

### Statistical analysis

R v.4.1.2 (51) was used for further analyses of data collected. For the description of *pedagogical suitability*, we proposed two items - I_17+I_18 (I_17 ‘I would use the web resource as it is even in standard teaching mode’, I_18 ‘I would use this activity as inspiration and modify it for standard teaching mode’) which we considered as complementary, i.e. negative correlation. We expected a split between web resources that are ready-to-use and resources that need more or less modification to be accepted by teachers as suitable. We tested this assumption with the Spearman correlation coefficient.

Principal component analysis (PCA) computed on the correlation matrix was used to describe the variability of the *pedagogical suitability* component. Data were not transformed as all items describing *pedagogical suitability* were collected in the same manner as 5-Point Likert scale (52). The decision on principal component count to be analysed was based on a scree plot of the explained variance of each of the principal components compared to the amount of variance each variable would contribute if all contributed the same amount.

*Interaction support* provided by the web resource was analyzed using a divisive hierarchical clustering. For clustering, we used the cluster package (53) with the similarity coefficient metric (54), which is suitable for categorical data (55); evaluation of the number of interpretable clusters was based on a scree plot of internal cluster variability and a silhouette index comparison (56).

A linear regression model to predict the overall score from individual subscales was fitted. We use a backward elimination to choose the final model.

To investigate the effect of subcategories from the *added value* group, we used the non-parametric Kruskal-Wallis test with Bonferroni correction on multiple testing.

All statistical tests were performed at the chosen significance level of 95%. Effect sizes were labeled following recommendations (57). 95% Confidence Intervals (CIs) and p-values were computed using a Wald t-distribution approximation.

## Results

We collected a total of 89 submissions representing 50 unique web resources (see S1 appendix for data table). A significant number of the resources were in national languages with local impact. Only two resources were mentioned more than three times. The resource most frequently mentioned (16 times) and the only one mentioned by teachers in at least three of the five countries was PhET (https://phet.colorado.edu) More than half (49, 55%) of the resources were mentioned only once. Teachers reported using nearly half of the activities even before corona (48%), 36% of the activities were newly discovered due to forced online teaching, and the remaining 16% were activities which teachers knew but did not use before corona. Most of the activities reported (82%) are free, 8% are available only after purchasing, or there are major differences between the free and paid versions, and 10% of the activities have minor differences between the paid and free versions. Most (76) of the resources implicitly had science content (e.g. WR_25 PhET, WR_5 periodic table of elements), 13 of the web resources could have any content depending on the teacher (e.g. WR_88 Mentimetr, WR_29 Kahoot, WR_85 Padlet).

### Digital elements

The teachers were asked to state a type of digital element for which they use the web resource. Most frequently, it was *DE_revision*, followed by *DE_simulation (e.g. population dynamics; input data, factors etc. and see results)* (Table 3). More than one digital element could be chosen and 3 was the most frequently used number of methods in their answer. The highest number of digital elements stated for one resource was 6.

**Table 3.** List of digital elements with the frequency among web resources. The percentage is counted from the number of resources (n = 89).

| Digital element type | Count [%] |
| --- | --- |
| DE_revision | 60 [67.4%] |
| DE_simulation | 40 [44.9%] |
| DE_investigation | 29 [32.6%] |
| DE_literature | 20 [22.5%] |
| DE_video | 20 [22.5%] |
| DE_experiment | 15 [16.9%] |
| DE_games | 11 [12.4%] |
| DE_virtual reality | 8 [9.0%] |

In the situation, where multiple answers were chosen at the same time, the combinations of *DE_revision* + *DE_simulation* or *DE_revision* + *DE_investigation* were the most common (N = 24), an overview of the combinations that occurred in more than 15 cases is given in Table 4.

**Table 4.** Most common pairs of digital elements reported together for a single web resource. The percentage is counted from the number of resources (n = 89).

| **Learning components couple** | **Count [%]** |
| --- | --- |
| DE_revision + DE_simulation | 26 [29.2%] |
| DE_revision + DE_investigation | 24 [27.0%] |
| DE_simulation + DE_investigation | 21 [23.6%] |
| DE_revision + DE_video | 16 [18%] |

### Pedagogical suitability

The highest sampled score of the *pedagogical suitability* was 35 (out of 35 possible), and the lowest *pedagogical suitability* score was 18 with the median value 27. Three most highly valued resources were all mentioning one website, PhET (WR_25, WR_70, WR_71). The two survey items - I_17+I_18 (I_17 ‘I would use the web resource as it is even in standard teaching mode’, I_18 ‘I would use this activity as inspiration and modify it for standard teaching mode’) - were designed to be complementary but turned out to be independent. The Spearman’s rank correlation rho between I_17 and I_18 is positive, statistically not significant, and very small (rho = 0.09, S = 0, p = 0.398).

The analysis of the primary components of *pedagogical suitability* showed the primary component PC1 explaining 37.8% of variability, the following axes 15.6% and 14.6% of variability, respectively, see Figure 1 and 2. Therefore, we interpret only the first component, graphically shown in Figure 2. PC1 is loaded with items I_17, I_19, I_20, I_21. That is “use as is”, “teach subject”, “structure clear”, “important ICT skill” see Table 5 for PCA loadings and item wording.

**Figure 1.**
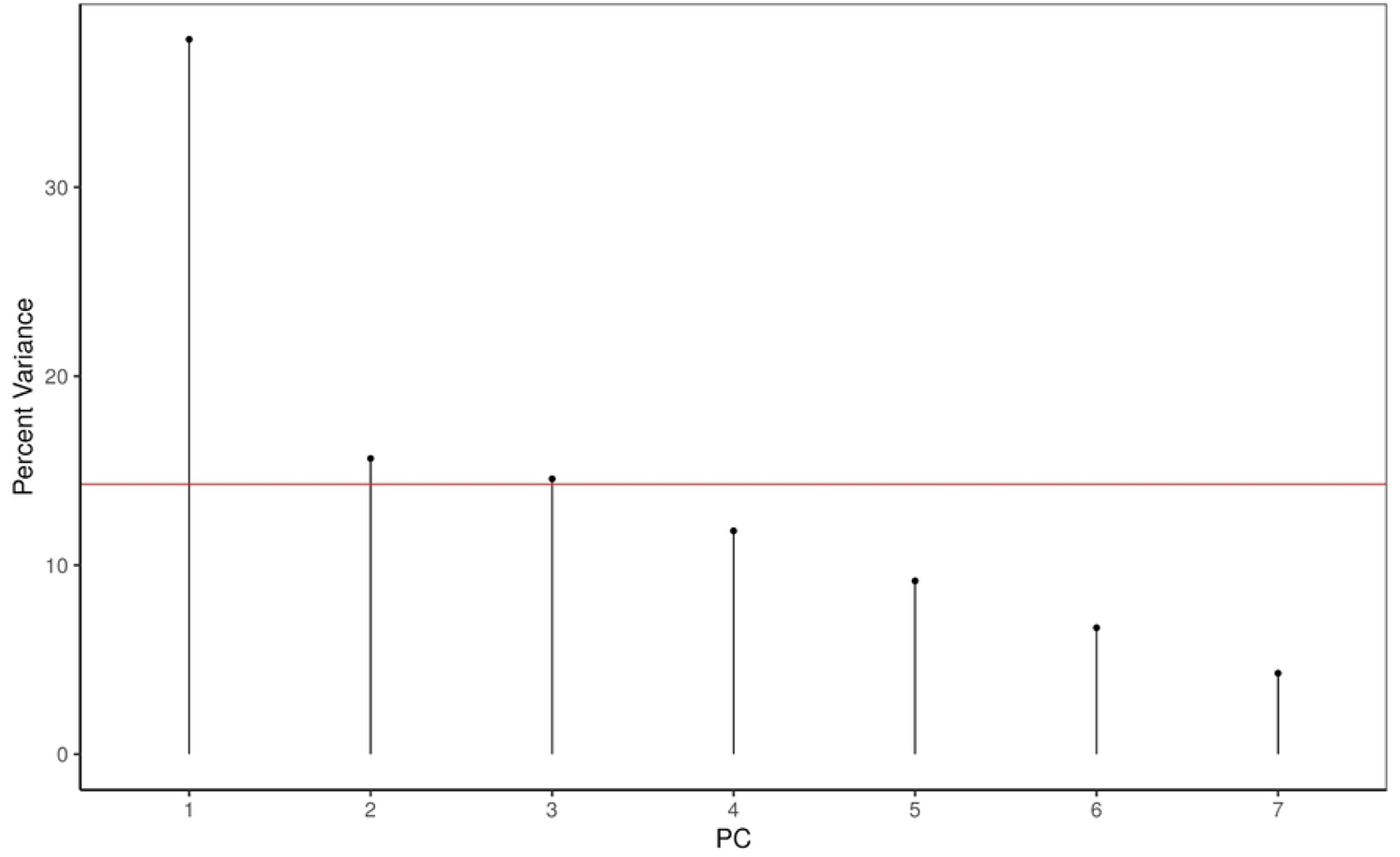
Explained variance percentage of primary components. The first primary component explains 38 % of variance. The red horizontal line indicates the amount of variance each variable would contribute if all contributed the same amount (14.3 %).

**Figure 2.**
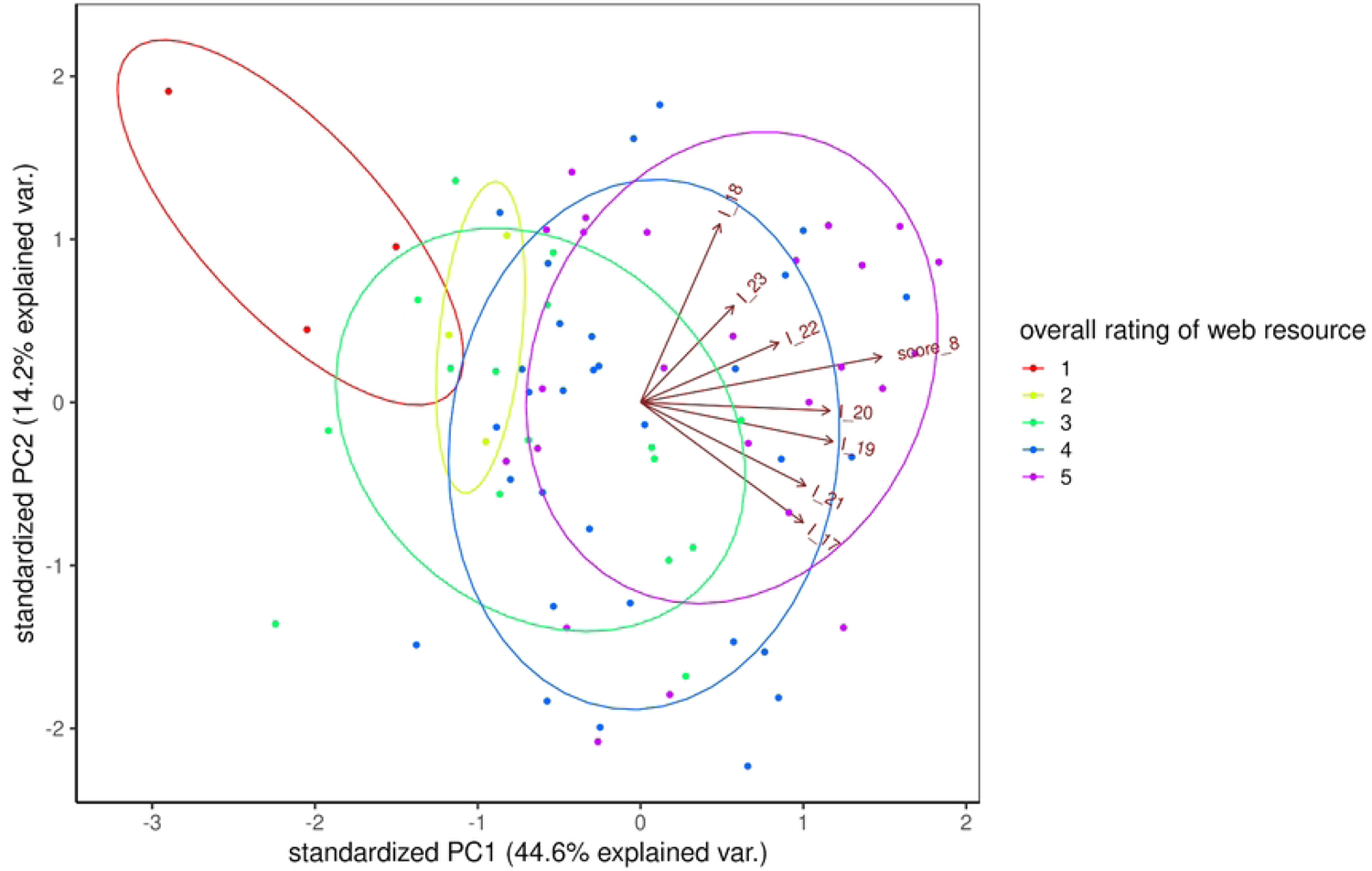
Visualization of the first and second primary components. The length of the arrows corresponds to the loadings of each item, the coloured ellipses highlight the distribution of the web resources with respect to their overall teacher rating. The trend of increasing overall rating of resources with respect to the positive direction of PC1 is clearly visible.

**Table 5.** PCA loadings in the first principal component. Loadings higher than 0.38 are highlighted in bold, as a theoretical cutoff if all variables contributed equally to that principal component.

| Item number | PC1 | Item wording |
| --- | --- | --- |
| I_17 | **0.4278753** | I would use this activity as it is even in standard teaching model |
| I_18 | 0.1373113 | I would use this activity as inspiration and modify it for standard teaching mode |
| I_19 | **0.5038127** | This activity helps to teach the subject |
| I_20 | **0.4891261** | The structure is clear |
| I_21 | **0.4074133** | It helps to teach an important ICT skill |
| I_22 | 0.3062552 | It helps to teach something else important (except subject topic and ICT) |
| I_23 | 0.2126473 | The visual is appealing |

Thus, the features forming PC1 are the main source of variability in the teachers’ evaluation of web resources (as overall ranking) in a linear model. The model’s intercept, corresponding to PC1 = 0, is at 4.02 (95% CI [3.85, 4.19], t(87) = 47.54, p < .001). Within this model, the effect of PC1 is statistically significant and positive (Std. beta = 0.61, 95% CI [0.44, 0.78], t(87) = 7.15, p < .001).

### Interaction support

The highest sampled score of the *interaction support* was 25 (out of 25 possible), and reached by three web resources (WR_42 Química en context, WR_70 PhET and WR_71 PhET), The lowest pedagogical suitability score was 5, median value was 17.

The *interaction support* category formed two distinctive clusters. The chosen number of clusters is supported by the Silhouette index, showing that two clusters are most dissimilar in between. Elbow plot does not indicate a clear cut-off and therefore we interpret two clusters (Figure 3). Group A was with interactions aiming inside school, student - teacher (I_24) and student - student (I_25) interactions which were supported by web resources like WR_88 Mentimetr, WR_29 Kahoot, WR_85 Padlet with which the activity is typically done synchronously. Group B missed the school interaction and could contain student’s interaction with society (I_26), nature (I_27), and family (I_28). Examples can be web resources WR_42 ‘Química en context’, or WR_17 ‘Climate facts’ which collects data on climate and climate change provided by scientific institutions (e.g. NASA, Eurostat, national data about weather and others) and compiles them into graphs and infographics for further use.

**Figure 3.**
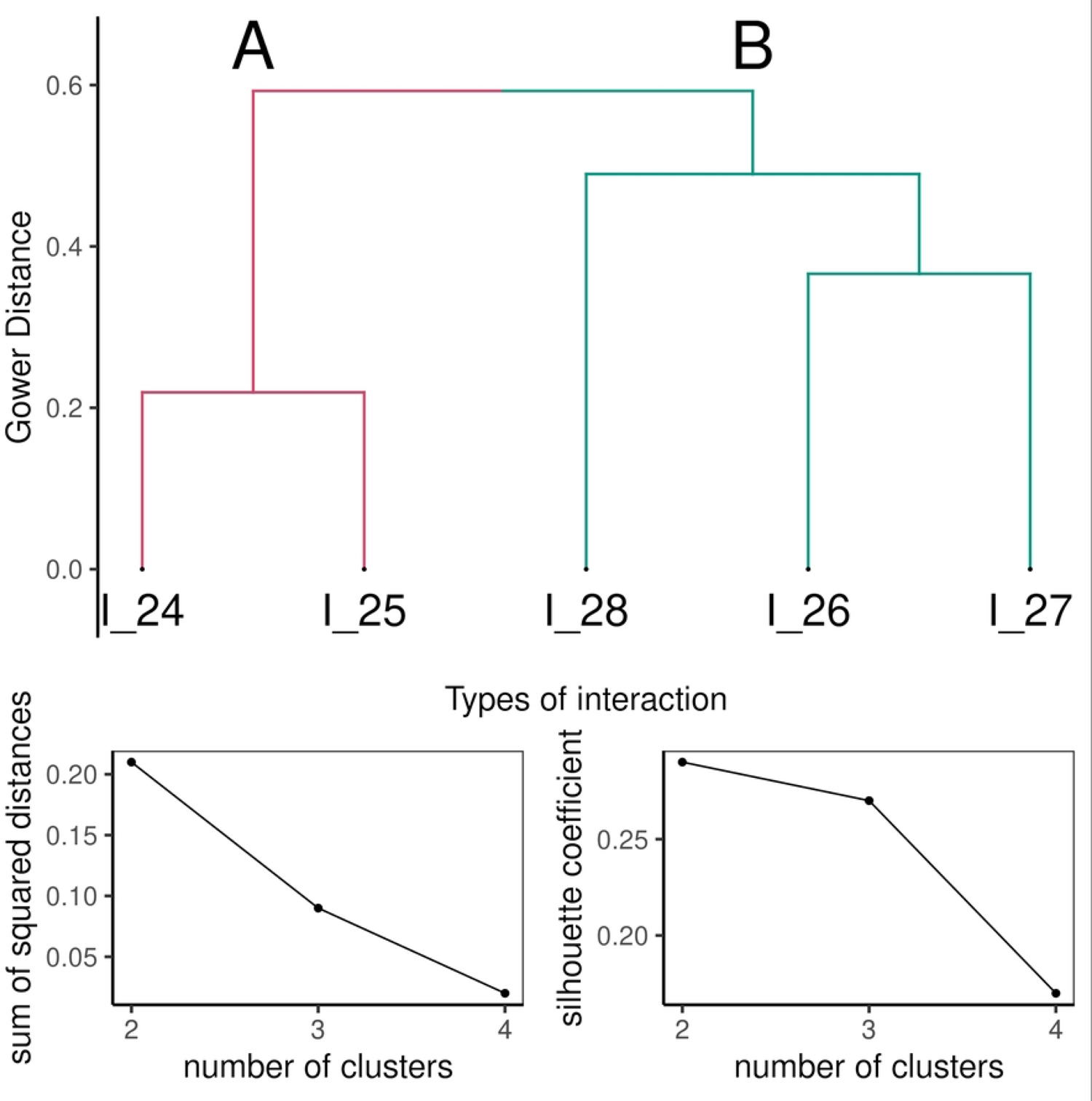
First row: Dendrogram of interaction support of web resources. Left cluster (marked A) represents interactions in school context; right cluster (marked B) represents interactions in the out-of-school context. Second row: Analysis of the elbow scree plot and silhouette index to select the number of interpreted clusters. The left plot represents the internal variability of clusters without sharp break. The right plot represents the Silhouette index, the dissimilarity of the clusters - the higher, the more dissimilar. The highest dissimilarity is just at two clusters.

The peer interaction (I_25) was the most frequently mentioned one (57 web resources, median = 4) followed by student-teacher interaction (I_24, 47 web resources, median = 4), student-society interaction (I_26, 41 web resources, median = 3) together with student-nature interaction (I_27, 40 web resources, median = 3). The least frequently mentioned interaction was the student-family one (I_28, 17 web resources, median = 3).

### Added value

The categories of web resource *added value* occurred with comparable frequency - I_35 (providing worksheets) was the least represented at 19 web resources (21%), and I_29 (supporting independent solving) and I_32 (safety) were the most frequently assigned - these were listed for 45 web resources (50.6%), with a median of 37.5.

More than a third, 32 web resources (36%), had no *added value* assigned. All categories were assigned to 11 web resources (12.4%). The remaining 46 web resources (51.7%) had between one and seven categories assigned (Figure 4).

**Figure 4.**
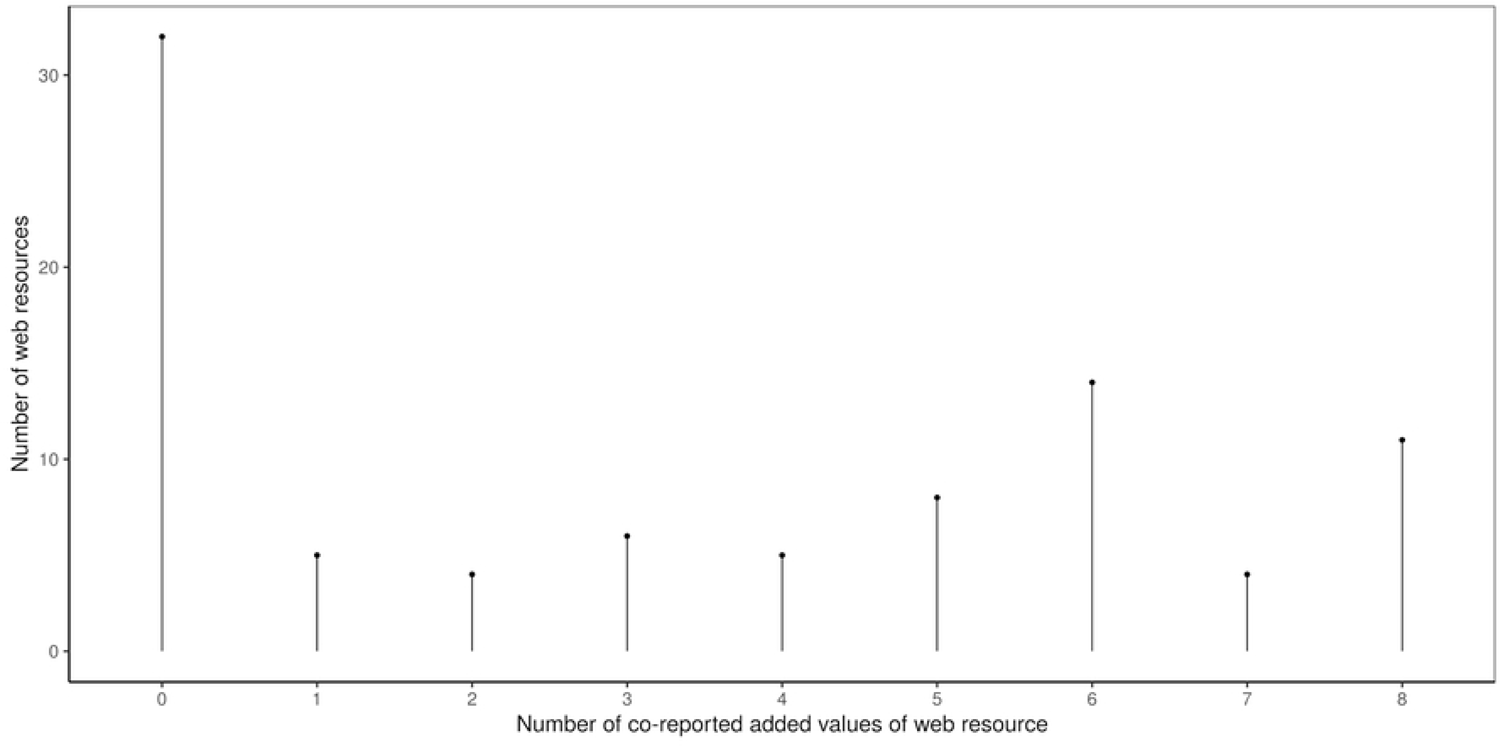
Number of added value categories listed together. More than a third of web resources had no added value assigned; see first bar - otherwise there was considerable variance in the number of *added value* categories listed together.

### Bloom’s taxonomy

Most (48, 53.7%) of the web resources were rated as “remembering”, aiming at low cognitive level.

### Overall rating

As a part of web resource reporting, the teachers gave an overall rating, a “number of points” to each web resource (the higher the better). The rating was mostly high (maximum 5 points of 5, median 4 points) as the teachers were asked only for the examples of good practice, the resources they wanted to recommend to others.

### Relationship between overall rating of web resource and individual subscales

We fitted a linear model to predict the overall rating of the web resource. As explanatory variables, we used *pedagogical suitability, interaction support*, *added value* and *Bloom’s digital taxonomy*, selecting the model by a backward selection. The accepted model formula is *overall rating* ∼ *pedagogical suitability* + *interaction support*. This model explains a statistically significant and substantial proportion of variance (R^2^ = 0.38, F(2, 86) =25.87, p < .001). Within this model:

- The effect of *pedagogical suitability* is statistically significant and positive (Std. beta = 0.44, 95% CI [0.22,0.66], t(86) = 4.02, p < .001)
- The effect of *interaction support* is statistically significant and positive (Std. beta = 0.23, 95% CI [0.02, 0.45], t(86) = 2.14, p = 0.035)

### Description of the most frequently mentioned web resources

*PhET* (phet.colorado.edu/) offers free interactive simulations for science and math that can be used in a variety of settings from demonstration experiments to individual exploration. Students test their hypotheses and answer questions by changing the setting of different variables and observing the results, and gain a deeper understanding of basic concepts. The simulations convey understanding of very small, quick, or fast processes which cannot be easily observed directly. The topics are, for example, the function of neuron, gene expression, color vision, natural selection, energy forms and changes, pH scale, diffusion, molecules and their polarity or shapes, atomic interactions or balancing chemical equations, solar system, electrical circuit, gravity force, density, vector addition, coulomb’s law, curve fitting, fractions, graphing quadratics, projectile motion and many more. Registered users gain access to teaching materials. The design of the simulations has been tested and proven effective (58–61). The web resource was mentioned 16 times in our survey, typically gained overall rating 5, and positive rating in *pedagogical suitability*. The evaluation of *interaction support* varied among the individual simulations evaluated as different teachers described different simulations or the web resource as a whole.

*Kahoot* (https://kahoot.com/) is suitable for those who want to create their own knowledge tests in a funny way. The user can be a teacher, a student or groups of students after creating an account. Kahoot can be used as a class competition when everyone gets the link at one time and also as a challenge for home repetition of the subject content. In this case, Kahoot is open for a limited time (one week) for everybody who gets the link. The students see their own position in the competition at the end of the game. The teacher has an overview of the difficulty of each question and also about placement of each player. There are more types of questions in the paid version than in the free version. The web resource was mentioned 4 times in our survey, typically gained overall rating 5 or 4, and would be used even in standard teaching mode, the visual was rated as appealing. The *interaction support* within school was mostly rated positive contrary to the out-of-school interactions. Kahoot represents an example of a web resource which can have any content, not only scientific one.

*Mozaweb (*https://www.mozaweb.com) is another paid application designed for science and humanities subjects. The registered teacher can use digital teachers’ books, presentations, e-learning, 3D animations, videos, and tasks to test students’ knowledge. The web resource was mentioned 3 times in our survey (overall ratings 5, 5, 1), and would be used even in standard teaching mode, the visual was rated as appealing. The *interaction support* was rated as neutral.

*‘Chemistry in context’* (Química en context, on Google sites) incorporates competency work at high school to support meaningful learning and motivate toward Chemistry through connections with everyday life. Among the activities there are many inquiry experimental ones, some of which take advantage of the potential of data recording and analysis equipment. In general, the activities focus on modelling, inquiry and argumentation (62) 2016). The web resource was mentioned 3 times in our survey (overall ratings 5, 5, 4), and would be used even in standard teaching mode, the structure was rated as clear. The *interaction support* within school was rated as neutral, and outside school (with nature, family and society) as positive.

‘*We know the facts*’ (https://www.umimefakta.cz) is a tool suitable for repeating or testing knowledge from almost all subjects. The school has to pay for the application to have the pupils in a class together and to allow the teacher to give them tasks and see the result. At the same time, individual students can use these activities individually for free; there is only a limit of 77 questions (approx. 30 minutes of revision) per day without registration. The tasks are done in a funny way at different difficulty levels. If teachers want to use the free version of the application, they can ask the students to send them a printscreen of the results. The web resource was mentioned 2 times in our survey (overall ratings 4 and 3), and would be used even in standard teaching mode. The *interaction support* within school was rated as neutral or nonexistent. The following two apps were mentioned only once, but we consider them worth sharing as a good practice.

The *geocaching* app (https://www.geocaching.com/sites/adventure-lab/en/) leads users outside. With the help of an application map, everybody sees the places which are near the position where they stand. Students (without the teacher) follow the path given by the teacher and collect the specified information about the places they visit. There are many prepared places with hidden content about their history or some interesting facts related to them, even in the free version. In the paid version, teachers can invent their own places with the content they need. The web resource can be both used as it is and modified for standard teaching mode. The *interaction support* among peers and with nature was rated as positive.

The virtual *microscope* app (https://www.ncbionetwork.org/iet/microscope/) is designed to familiarize the student with the microscope in advance. Everybody with the link gets into the application for free to see the structure of the microscope and can explore selected plant, animal, human and bacteria microscopic slides. The slides are also described, and the user can change magnification, adjust light, and both coarse and fine focus. The application points out that the magnification needs to be changed gradually or shows a broken slide if such a situation should happen in reality. There is also a test on care and usage of the microscope, calculating magnification, and terminology. The *pedagogical suitability* was rated positively, the *interaction support* as neutral.

## Discussion

We identified web resources used and recommended by science teachers in five EU countries for teaching science practicals. To recall our conceptualization, a *web resource* is an objectively existing online resource that teachers incorporate into their teaching in a (very) variable way, bringing a digital element into their practice. Among the reported web resources, there were a few that were repetitive across countries (PhET, Interactive Simulations for Science and Math, with many language mutations, Kahoot in which teachers add content in their language in two, and three web resources were mentioned in Czechia and Slovakia where there is little, if any, language barrier), but a large proportion of these were unique to countries. This situation highlights the need for an international design of circumstantial studies and raises two possible interpretations. Either the local resources are limited in their international impact by the language barrier (and the resulting low speed of dissemination into the global space), or the resources have some specificity to a given curriculum design or school tradition. This would be very interesting to find out, for example, in relation to cultural dimensions (63). The teachers started using approximately half of the reported web resources due to the forced distant teaching. This shows that the respondents were open to including new resources and made them part of science education. They have undergone the two critical phases necessary for adopting teaching and learning based on computing devices formulated by (1) i) introduction and gaining practice, and ii) implementing the tools into lessons. It confirms the data collection timing was well chosen for such a type of study.

The consequent analysis allowed us to interpret shared characteristics of the web resources and discuss to what extent the theoretical frameworks (9,10,24) coincide with lived teaching practice. We have used only some keystones from the frameworks which were relevant to our aim: characterization of the web resources used for online science practicals teaching. From these, we point to *interactions* building relationships and community out as a set of items on its own. The other characteristics (supporting independent solving, active learning and creativity, difficult differentiation, safety, authenticity, flexibility of topics) formed an *added value* category together with items relevant to science practicals ‘providing worksheets’ (Klein et al., 2021) and ‘connecting different organizational levels’ (48, 49).

The vast majority of web resources had multiple digital elements mentioned, even ones that were not entirely expected on first impression of the content of the resource. This helped us to understand that the degree of variability with which teachers conceptualize the incorporation of a given resource, i.e. the nature of how they incorporate its use into their teaching and what kind of ‘digital element’ they create from it, is enormous and we estimate that it exceeds the expectations of the creators of the resources - the creativity of the creators of web resources is added to the creativity of the educators who push their work into sometimes unexpected positions. For this study, this creates a source of bias because we did not expect such diversity and we took the web resource as the primary entity of interest. Hence, it is not possible to determine whether, for example, added value refers to all types of use of a given resource, or whether teachers associated it with some particular specification of the digital element. We interpret the data in accordance with the data collection concept with this potential bias in mind. The most frequently mentioned web resource was PhET which also represented many of the most frequent characteristics; therefore, we use it as an example when discussing the findings of this study below.

The most frequently mentioned *digital element* was knowledge *revision*. This may imply the prevailing ‘remembering’ level of Bloom’s taxonomy although the most frequently mentioned digital elements, *revision* and *simulation*, also formed the most frequently mentioned digital elements couple as in the example of PhET. And revision by using simulations suggests that in reality, the teachers aimed at higher cognitive levels. The level of Bloom’s taxonomy was coded by the authors (only as binary ‘remembering’ – ‘higher’) as we believe it helps better understand the role in teaching practicals. However, in this arrangement, interpretation was difficult and sometimes even ambiguous because web resources could be used in different ways.

The PhET web resource also had the characteristics important for the overall rating included in *pedagogical suitability*: ready to use without modifications to teach subject content and ICT, clear structure. Consequently, it was the best rated web resource within *pedagogical suitability*. This is in concordance with the literature as PhET has been demonstrated to be an effective learning tool and it is gratifying that teachers across Europe are incorporating it into their (online science practicals) teaching (60,61,64).

As (social)interactions of different types are seen as one of the keystones of online education (9, 10) we used them as a separate set of items and also added student-nature interaction which is fundamental for science education. Our results show that teachers most recommended apps that promoted interaction between students, student and teacher, student and nature, and student and society. We hypothesize that teachers did not consider student+family interaction as important, or instead purposely encouraged interactions outside the family during lockdown, as it could be assumed that most of the students spent the majority of the lockdown time with their families. Interaction with peers is very important and it was challenging to maintain it during lockdowns (65); therefore, it is good news that the teachers recommended the web resources encouraging student+student interaction. Student+teacher interaction, from which especially individual feedback is highlighted (66), was also supported by most of the used web resources. Students’ interaction with nature is essential for both physical (67) and mental health (68) and should be one of the goals of science education. As such, it was seen as important when choosing web resources for online teaching. In the case of PhET, the *interaction support* evaluation differed among teachers, as the web resource offers a wide range of different simulations and the specific work assignment may vary.

As with most web resources, PhET is available for free and was used also before COVID-19 forced online teaching. Contrary to most of the web resources, PhET simulations represent many of the *added values*. Surprisingly, 36% of the recommended web resources did not provide any *added value,* and these characteristics based on research and literature recommendations (9,24,25) did not influence the overall rating of the web resources and were not in the regression model. Teachers evidently also used web resources, which according to theory did not meet the requirements for online teaching of science practicals. The question is whether they were aware of this and used web resources to supplement other activities, or whether this was the result of forced online teaching that was not well thought out, lacked integration, and thus was unlikely to be effective (7, 8). There remains also the option that the survey items were misunderstood.

Based on the proposed model, the overall rating of the web resources was influenced neither by Bloom’s taxonomy nor by *added values* derived from (9,10,24). Online teaching practice seems to be different from what the theory assumes and demands if teachers still use the web resources as a part of effective online science teaching (7, 8). On the other hand, it is possible that the usage of the web resources is not that well thought through and is a result of rather random choices and more integration is needed. The model shows that the *pedagogical support* and *interaction support* influence the overall rating. This seems to be straightforward, as the pedagogical support in addition to others describes how well the web resource prepared for direct usage in science education and the interactions are fundamental to education.

### Limitations of the study

As in other similar studies, only teachers willing to participate in the survey shared their ideas with us. Not all science teachers in the countries were asked to participate, rather each institution used its own communication means to contact science teachers, which creates a bias. We only collected the web resources the science teachers would recommend; consequently we don’t know which web resources they have tried and possibly rejected. Therefore, we cannot distinguish which ones they don’t want to use and which ones they don’t know. As mentioned above, a web resource is taken as a base entity in our survey, and therefore is impossible to distinguish reported web resource features to the specific digital element context.

## Conclusion

We built on previous work on the typology and description of the roles of digital elements in teaching and specified features that are essential for teachers’ decision-making when integrating a given resource into teaching. By analyzing examples of good practice from five European countries, we found that teachers preferred web resources that provided knowledge revision and virtual simulations, could be used without modification to teach the content and ICT, and promoted social and nature interactions. Pedagogical suitability (e.g. teaching content, ICT, clear structure) and support of interactions were the most influential factors in teachers’ rating. Surprisingly, characteristics considered essential by educational research experts, such as evaluation according to Bloom’s taxonomy, added value (e.g. linking organizational levels, supporting independent solving, creativity, flexibility, allowing differentiation of difficulty levels, providing workshops) did not influence the overall rating of the web resources. Thus, we point to the gap between theory and practice.

## Acknowledgement

This work was supported by Erasmus+ project My Home - My Science Lab. We are grateful to the members of My Home - My Science Lab project and teachers involved in the study.

